# Patching Science – amending the literature through version control

**DOI:** 10.1101/2022.04.13.487348

**Authors:** Adam Kane, Bawan Amin

**Author notes:** Corresponding author –.

## Abstract

The ideal of self-correction in science is not well served by the current culture and system surrounding amendments to published literature. Here we report on a survey (N = 132) that highlights academics’ dissatisfaction with the status quo and their support for an alternative approach. We then describe our view of how amendments could and should work by drawing on the idea of an author-led version control system. Here authors would include a link in their published manuscripts to an updatable website (e.g. a GitHub repository or similar) that could be disseminated in the event of any amendment. Such a system is already in place for computer code and, as such, requires nothing but buy-in from the scientific community - a community that is already evolving towards various open science frameworks. This would remove a number of frictions that discourage amendments thus leading to an improved scientific literature and a healthier academic climate.

## The problem

Science is held to lofty ideals. Chief among them is its supposed capacity for self-correction. However, we, and many other academics, argue that the current model of scientific publication hinders this capacity (Barbour et al. 2017; Teixeira da Silva and Al-Khatib 2021; Allison et al. 2016). People make mistakes and the peer review system can’t catch them all (Molckovsky, Vickers, and Tang 2011; Kendall et al. 2019). Yet the number of steps involved in correcting or supplementing previously published research acts to discourage what should be a straightforward process that is in the hands of the original authors (Barbour et al. 2017; Teixeira da Silva and Al-Khatib 2021). After all, it is the authors who are best placed to correct their own work.

This is to say nothing about academia’s difficult relationship with authors owning mistakes (Barbour et al. 2017; Teixeira da Silva and Al-Khatib 2021; Rohrer et al. 2021). Indeed, an article in Times Higher Education (2017) explored the “emotional, reputational and practical” barriers standing in the way of correction (Else 2017). The same article showed a significant proportion of those surveyed would not report a major mistake in a high impact paper that affected the paper’s conclusions. Certainly, nobody likes to be wrong and the mental anguish of people who go through correcting or retracting something they spent years of work on is palpable (Chawla 2019; Conroy 2020). But, that some commentators deem these acts as heroic shows just how far away we’ve moved from the self-correcting ideal– heroes are exceptional, mistakes are not (Vuong 2020).

Then there is the simple opportunity cost of dealing with relatively minor issues in a paper. Those that vex but aren’t worth the time and effort to correct (Else 2017). Or instances where you have a new dataset that could supplement previous work but isn’t worth publishing separately.

This is all particularly aggravating because the internet, especially Web 2.0, offers us a way forward through version control. The static face of academic publications is a holdover from times where print held sway, but the dynamic and reflective nature of the internet is much more in keeping with how science should operate (O’Dea et al. 2021). We propose that researchers should create versions of their own papers that record amendments to their work while retaining the original published manuscript for reference. Here we report results from our own survey to assess the level of support for our proposal, detail how the system would work including its many benefits, and note some responses to potential counterarguments.

## Survey

We developed a survey to first inquire about scientists’ knowledge, exposure, and experience with the process of issuing a correction (see supplement). We also asked the following statement that relates directly to our proposed solution:

*“Imagine adding a link to published papers, which would direct readers to an online, open and updateable repository (e*.*g. GitHub, OSF, etc*.*). Such a framework would be used by the authors of the paper to add any update to said paper. Updates can be amendments, text corrections, additional data and analysis. These updates would not alter the journal’s version of record. This system would not need any involvement from the journals (except providing the link)*.*”*

We then finished with a suggestions box so that people could expand on this or related points. We built the survey using google forms and disseminated it over Twitter as well as contacting 13 researchers who we know had an interest in this area with the request to spread it through their Twitter network.

## Results

We recorded 132 respondents (see supplementary material for full results), two thirds of whom were in the life sciences across career stages. Remarkably, almost a third of respondents were unaware it was possible to make amendments to peer-reviewed publications. In keeping with previous results, the percentage of people who have considered amending their work exceeded the percentage of those who have attempted an amendment. Out of the 19 people who successfully amended their work, only a quarter (5) agreed that the current system worked well, despite their previous success. The remainder were either neutral (8) or disagreed (6). There were a variety of reasons given by the 34 people who hadn’t formalized an attempt for amendment including lack of clarity as to how to proceed (20), hassle with the publishers (12), the time needed (9), embarrassment (8), and scorn of their peers (3). In response to our proposed solution only around 7% (9) of people disagreed with the idea, the rest were in support (∼61%, 81 people) or conditionally so (∼32%, 42 people).

## Proposal

Here we further flesh out our proposal beyond the statement from the survey. We suggest that every published paper includes a link to a page controlled by the authors, for instance a GitHub repository (or equivalent e.g. Open Science Framework). This is already in place for things like supplementary data and code in line with open science practices (O’Dea et al. 2021; Sandve et al. 2013). Here, the authors could detail errors, amendments, and additional analyses using the ‘readme’. This readme would include extensive information on the differences between the versions. We note that these need not be reserved for coding issues, everything from a fundamental coding error to an awkward sentence could be fixed. Because of the version control features inherent to Git, the original, peer-reviewed version would still be accessible. Further, many researchers already use GitHub to record new versions of R packages commonly used by scientists across many fields (e.g., Hadley Wickham’s tidyverse – https://github.com/tidyverse/tidyverse/releases). Authors could draw attention to this new version by creating a new file on their Google Scholar profiles, ResearchGate accounts etc. We suggest a simple X, Y, Z numbering system that follows best practice for version control elsewhere through semantic versioning (https://semver.org/) where X is a major additional analysis, Y is a minor additional analysis and Z is a correction. This would display as – ‘Darwin, C. (1859). On the Origin of Species. - VERSION 1.0.0’ for the initial publication. And a correction as ‘Darwin, C. (1859). On the Origin of Species. - VERSION 1.0.1’ with updates containing a link to the GitHub release page (see Fig. 1).

**Fig.1.**
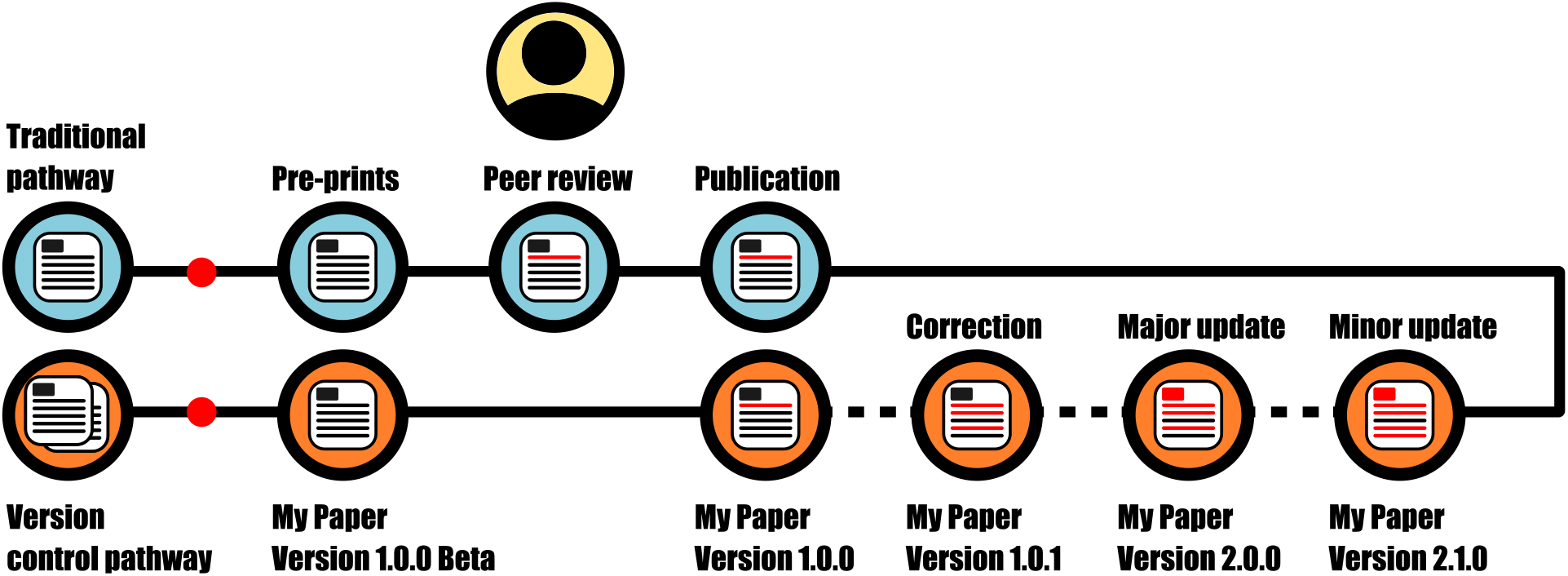
Graphic representation of our proposed system that shows its relationship with the current system.

## Benefits

We believe there are multiple benefits to our proposed system. The number one benefit is that it allows previously published work to be amended and/or updated in a straightforward way because it is author led. This takes out the middleman of the journals who do not need to be involved at any step once the paper is published in its initial, peer-reviewed state (see responses to Question 9 in the supplement). We suspect that this relatively frictionless system would engender a culture of correction including an awareness of correction as a possibility (Teixeira da Silva and Al-Khatib 2021). Adhering to semantic versioning would allow authors to have a natural flow if they use preprints by referring to this version as the beta stage. Further, it would offer plenty of grist for students to verify/update/correct the work of their supervisor – typically a principal investigator who may have more time constraints. This would stand to benefit the students who could point to tangible outputs from their own efforts on their CVs. It would also empower authors to raise flags about their own work even if they haven’t yet had time to address them (cf. the Loss-of-Confidence Project (Rohrer et al. 2021)). Historical works that do not have an internal link to GitHub could still be updated using this system by establishing a page for the paper post hoc.

We argue that the concerns raised by people in our survey as well as our own thoughts of potential shortcomings are not unique to this system. Disagreement among co-authors, visibility of the amendment and malpractice are some of the issues raised that pertain just as much to the current system of corrections (Molckovsky, Vickers, and Tang 2011). Indeed, we note that ours is an additional system, it allows for immediate corrections, where authors can still go through the established route if they desire.

## Conclusions

Our findings add more weight to the view that the current system of academic publication hinders science by disincentivising corrections and amendments to published literature (Allison et al. 2016). A system of version control, as we detail here, would offer numerous benefits to what is an inherently dynamic and imperfect process of discovery (Ioannidis 2005), all the while keeping the version of record. The fundamentals to this system are already out there and simply require buy-in from the community. Although it is not a perfect system, we believe a constantly updating and updateable literature better reflects the ideal of science. With this, we can take another step towards a more robust and reliable scientific literature and an improved working academic climate.

## Ethics

Ethical clearance was granted by UCD’s Human Research Ethics Committee - Research Ethics Reference Number: LS-E-21-281-Amin-Kane.

## Acknowledgements

Thanks to the Laboratory of Wildlife Ecology and Behaviour at UCD for testing our survey and providing valuable feedback on the questions.

## Open Science statement

Updates of this paper can be found following this link: https://github.com/kanead/Corrections

