## Supplementary material for "Patching Science – amending the literature through version control": SurveyResults.html

Survey Results


### Survey Results

### Description

To explore how peers experience the current situation regarding amendments of scientific papers, we ran a survey titled “Amending the scientific literature: A survey”.

The survey was distributed mainly via Twitter for the first time on January 10th, and remained open for a period of three weeks until January 31st. During this period, we received a total of 132 individual responses.

In this file, we present the results of the survey. We will go through each question and give a breakdown of the responses.

##### Question 1: What field are you currently in?

Mandatory question, only one answer possible. Various options provided. Participants were allowed to give a unique response. To maintain a good overview, we collated unique answers, i.e. answers with a frequency of 1, into a new category named “Other”.

Although we had participants from multiple disciplines, we see that the majority are active in Life Sciences, namely 65% (86/132).

##### Question 2: What is your position?

Mandatory question, only one answer possible. Participants were allowed to give a unique response. To maintain a good overview, we collated answers with a frequency lower than 5, into a new category named “Other”.

We can see that our survey has a good representation from different career stages.

##### Question 3: What is your gender?

Mandatory question, only one answer possible. Participants were allowed to give a unique response.

We can see that we had a pretty much 50/50 split in genders, with n = 3 participants indicating that they prefered not to specify.

##### Question 4: Have you published in a peer-reviewed journal before?

Mandatory question, only one answer possible. Participants were only given “Yes” or “No” as options.

We can see that more than 90% of our participants have published in a peer-reviewed journal.

##### Question 5: Are you aware it is possible to make amendments to peer-reviewed publications?

Mandatory question, only one answer possible. Participants were only given “Yes” or “No” as options. Since we think that the answers may be influenced by a participant’s experience with publishing, we show two plots here. The first one is of all participants. The second one only includes participants who had indicated that they have published before.

We can see that, surprisingly, the proportions show little effect of previous experience. In both cases, even when only including individuals that have at least one peer-reviewed paper, roughly 30% is not aware that papers can be amended.

##### Question 6: Have you ever considered amending some (minor or major) aspect of your previously published work?

Mandatory question, only one answer possible. Participants were only given “Yes”, “No” or “I have not previously published” as options. Since the question relates to individuals that have published, we have here omitted the responses (n=11) that indicated “I have not previously published”.

We can see that roughly a third of the participants has considered amending previously published work.

##### Question 7: Have you ever attempted to amend an error in one of your published works via the journal, and were you successful?

Mandatory question, only one answer possible. Participants were only given “Succesful attempt”, “Unsuccesful attempt” or “Never made an attempt” as options. To break down the results, we will show two figures again. The first one shows the responses of all participants. The second one shows only the responses of participants that have made an attempt.

We can see that roughly 20% (28/132) of the participants have previously made an attempt to amend some of their work. Out of these 28 participants, 68% have made a successful attempt and 32% an unsuccesful attempt.

##### Question 8: If your attempt was successful: would you agree that the current amendment system works well?

Optional question, multiple answers possible. Participants were given various options and also the option to formulate a different answer. Here we only include the individuals that have previously answered that they have made a successful attempt (n = 19)

Despite having made at least one succesful attempt, only a quarter of the participants indicated that they think the current system works well (Agree or Strongly agree; 5/19; 26%). The largest group of participants didn’t agree or disagree with the statement (Neutral; 8/19; 42%), followed by the group of participants that think that the current system does not function well (Disagree or Strongly disagree; 6/19; 32%).

##### Question 9: If the attempt for amendment was unsuccessful: Why?

Optional question, multiple answers possible. Participants were given various options and also the option to formulate a different answer. Here we exclude individuals who responded “I have made no attempts”. That, together with the fact that this question was optional, led to a sample of n = 11 individuals.

Participants reported “Editors/Journals not being cooperative” as the main reason for why amendments were unsuccessful.

##### Question 10: If you haven’t formalized an attempt for amendment: Why not?

Optional question, multiple answers possible. Participants were given various options and also the option to formulate a different answer. After omitting rows of participants that answered “I have not considered an attempt”, the remaining dataset consisted of n = 34 participants.

Compared to Question 9, we see that participants indicate that the main reason for not formalizing attempts is because they find it unclear how to proceed (58%). In addition, we see again that the confrontation with editors/journals is something that can withhold individuals from making amendments, since 35% gave this as a reason.

#### Our proposal and the final question of the survey

In our main manuscript, we make a case for implementing an additional amendment system that can facilitate the whole process of amendments of scientific papers within science as a whole. We wanted to explore whether there would be support for our proposal, and therefore we asked the participants of our survey whether they would welcome our suggestion.

Specifically, we presented them with the following framework:

###### ***Imagine adding a link to published papers, which would direct readers to an online, open and updateable repository (e.g. Github, OSF, etc.). Such a framework would be used by the authors of the paper to add any update to said paper. Updates can be amendments, text corrections, additional data and analysis. These updates would not alter the journal’s version of record. This system would not need any involvement from the journals (except providing the link).***

##### Question 11: Would you welcome such a framework for published papers (ran by authors)?

Mandatory question, only one answer possible. Participants were given “Yes”, “No” and “Maybe” as option, together with the additional option to formulate their own answer answer. Since all unique answers included some sort of conditional support, we decided to group all of these, together with every “Maybe” answer, into one category named “Conditional support”.

We found that the majority of the participants are in support of our proposed framework (81/132; 61%). Another 32% (42/132) indicated conditional support, and only 7% (9/132) were against our proposed framework.
